# Advances in mixed cell deconvolution enable quantification of cell types in spatially-resolved gene expression data

**DOI:** 10.1101/2020.08.04.235168

**Authors:** Patrick Danaher, Youngmi Kim, Brenn Nelson, Maddy Griswold, Zhi Yang, Erin Piazza, Joseph M. Beechem

## Abstract

We introduce SpatialDecon, an algorithm for quantifying cell populations defined by single cell RNA sequencing within the regions of spatially-resolved gene expression studies. It obtains cell abundance estimates that are spatially-resolved, granular, and paired with highly multiplexed gene expression data.

SpatialDecon incorporates several advancements in the field of gene expression deconvolution. We propose an algorithm based in log-normal regression, attaining sometimes dramatic performance improvements over classical least-squares methods. We compile cell profile matrices for 27 tissue types. We identify genes whose minimal expression by cancer cells makes them suitable for immune deconvolution in tumors. And we provide a lung tumor dataset for benchmarking immune deconvolution methods.

In a lung tumor GeoMx DSP experiment, we observe a spatially heterogeneous immune response in intricate detail and identify 7 distinct phenotypes of the localized immune response. We then demonstrate how cell abundance estimates give crucial context for interpreting gene expression results.

## Introduction

Single-cell RNA sequencing defines the cell populations present within a tissue. But this catalog of cell types begs a question that scRNA-seq cannot answer: how are these cell types arranged within tissues? Spatial gene expression technologies^1,2^, measure gene expression within minute regions of a tissue, but do not report abundance of cell types within these regions.

Here we introduce SpatialDecon, a method to quantify cell populations within the regions of spatially-resolved gene expression studies. SpatialDecon obtains cell type abundance measurements that discriminate closely related cell populations and are paired with expression levels of hundreds to thousands of genes. These measurements reveal the spatial organization of cell types defined by scRNA-seq, and they give context to gene-level results, resolving whether a gene’s expression pattern reflects differential expression within a cell type or merely differences in cell type abundance.

Our method employs gene expression deconvolution, a class of algorithms designed to estimate the abundance of cell populations in bulk gene expression data. Figure 1 summarizes the workflow of these algorithms. Given the inputs of gene expression data and pre-specified cell type expression profiles, deconvolution algorithms seek the cell abundances that best fit the data. The deconvolution field is still developing, and existing algorithms developed for use in bulk expression data^3,4,5^ are not optimized for spatial gene expression platforms. We have developed algorithms and data resources to make deconvolution more robust and widely-applicable. These advances enable accurate deconvolution in spatially-resolved expression data.

**Figure 1:**
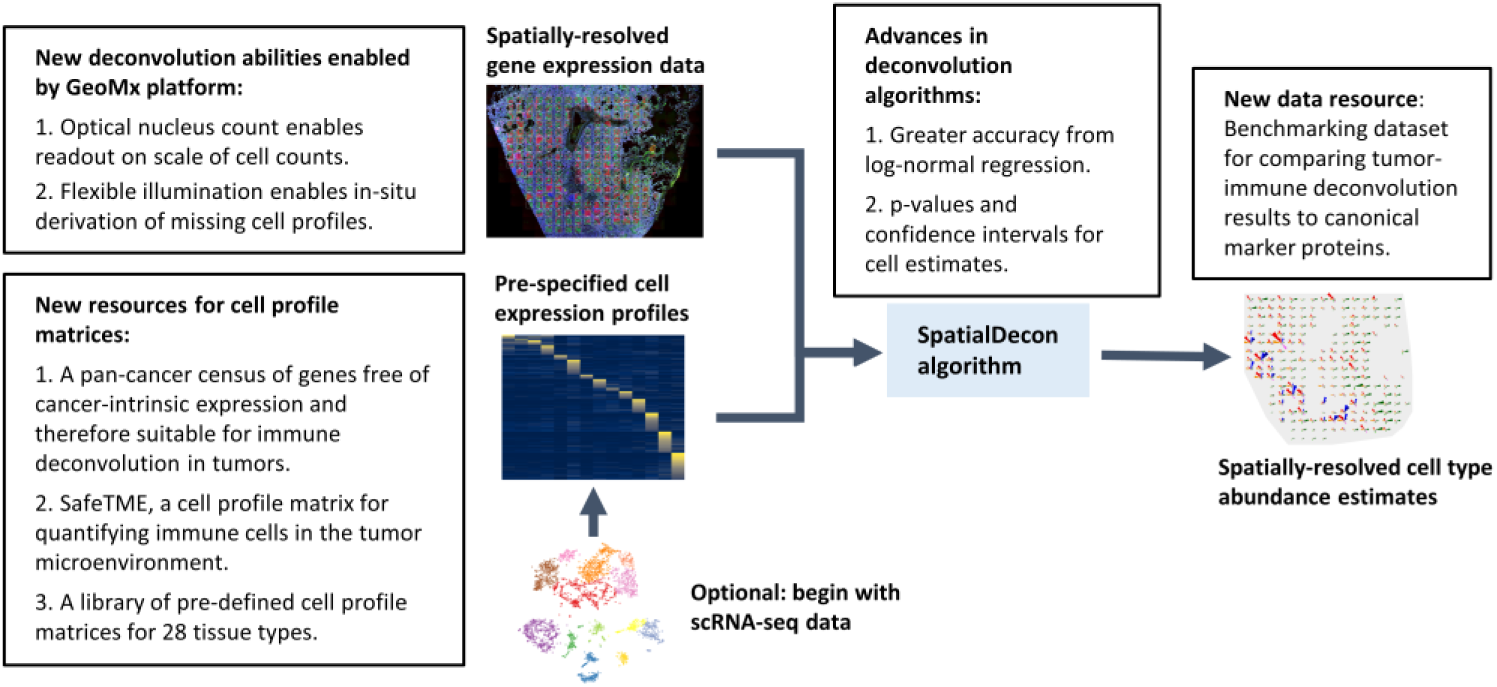
Overview of algorithm and advancements to the deconvolution field. The image summarizes the deconvolution workflow. Text boxes summarize new developments described in this manuscript.

## Results

### A novel algorithm achieves accurate deconvolution in spatially-resolved gene expression data

Gene expression data has extreme skewness and inconsistent variance, but most existing deconvolution algorithms are based in least-squares regression and implicitly assume unskewed data with constant variance^3,4,5^. The skewness and unequal variance of gene expression are corrected by log-transformation (Supplementary Figure 1). We therefore propose to replace the least-squares regression at the heart of classical deconvolution with log-normal regression^6^. This approach retains the mean model of least-squares regression while modelling variability on the log-scale. SpatialDecon, the algorithm implementing this procedure, is described in the Online Methods.

To evaluate the performance of log-normal vs. least-squares deconvolution, two cell lines, HEK293T and CCRF-CEM (Acepix Biosciences, Inc.), were mixed in varying proportions, and aliquoted into a FFPE cell pellet array. Expression of 1414 genes in 700 µm diameter circular regions from the cell pellets were measured with the GeoMx platform.

Three deconvolution methods were run: non-negative least squares (NNLS), v-support vector regression (v-SVR), and constrained log-normal regression (Algorithm 1 in Online Methods). Four gene subsets were used: the most informative genes, the least informative genes, all genes, and the genes with low-to-moderate expression (Fig 2a).

**Figure 2:**
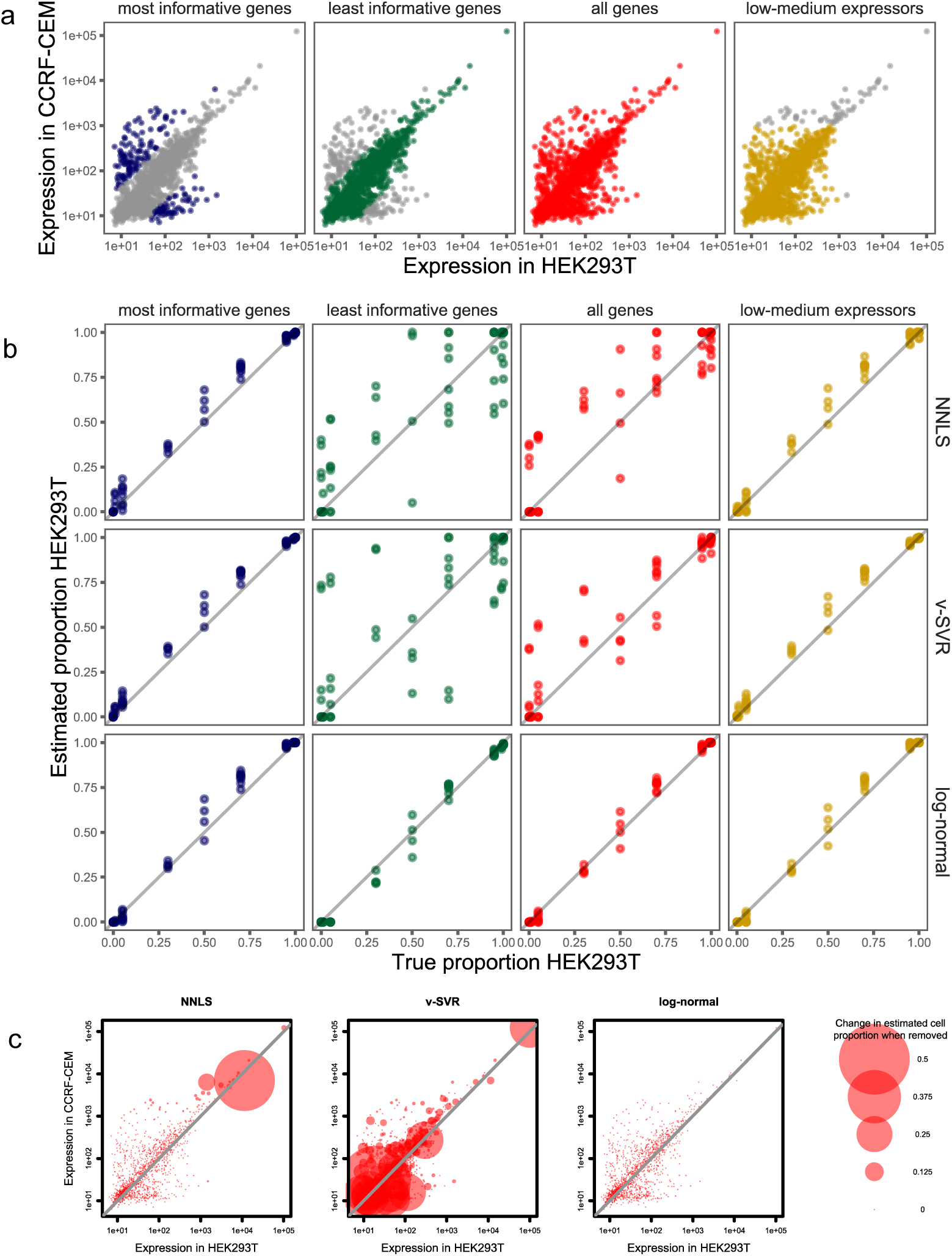
Comparison of deconvolution algorithms in mixtures of two cell lines. The cell lines HEK293T and CCRF-CEM were mixed in varying proportions, the GeoMx platform was used to profile each mixture’s gene expression, and 3 deconvolution methods were used to estimate the cell lines’ mixing proportions from the gene expression data. **a**. Expression profiles of the two cell lines. Colors denote subsets of genes used in separate deconvolution runs. **b**. Accuracy of deconvolution algorithms. Horizontal position shows each sample’s true proportion of HEK293T; vertical position shows estimated proportion. Each column of panels shows results from a single gene set; each row of panels shows results from a single deconvolution algorithm. **c**. Influence of each gene on the deconvolution result from a single mixed sample. Point size shows how much removing each gene changes the estimated mixing proportion. Each panel reports genes’ influence under a different deconvolution algorithm.

Deconvolution accuracy was evaluated by comparing the cell lines’ estimated vs. true mixing proportions. Log-normal deconvolution was accurate in all gene sets, with estimated proportions always differing less than 0.19 from true proportions (Fig 2b). In contrast, the least-squares-based algorithms (NNLS and v-SVR) failed in the “least informative” and “all genes” gene sets, with estimated cell proportions differing from true proportions by as much as 0.50 for NNLS and 0.73 for v-SVR. The gene sets in which least-squares methods failed are distinguished by the presence of high-expression genes.

### Classical deconvolution methods based in least-squares regression assign excessive influence to small subsets of genes

To investigate the poor performance of least-squares-based methods, we measured the influence of each gene on deconvolution results from a single cell pellet with an equal mix of HEK293T and CCRF-CEM. Each gene’s influence was measured as the difference in estimated HEK293T proportion using the complete gene set vs. a leave-one-out set omitting the gene in question.

The least-squares methods NNLS and v-SVR both had genes with excessive influence on deconvolution results, while the log-normal method was not subject to outsize influence from any genes (fig 2c). For NNLS, a single high-expression gene changed the model’s estimated mixing proportion from 66% to 22%, a remarkable impact on a fit derived from 1414 genes. In v-SVR, genes across all expression levels showed excessive influence; the most influential gene changed the estimated proportion from 31% to 83%. Removing the highest-influence gene from the log-normal deconvolution changed the estimate from 54.6% to 54.3%.

### Pan-cancer screen for genes with negligible expression in cancer cells identifies genes safe for immune deconvolution in tumors

Deconvolution of immune cells in tumors encounters another complication: genes expressed by cancer cells contaminate the data, causing over-estimation of the immune populations also expressing those genes. We analyzed 10,377 TCGA samples to identify a list of genes with minimal contaminating expression by cancer cells. We used marker genes^7,8^ (Supplementary Table 1) to score abundance of immune and stromal cell populations in each sample, and we modelled each gene as a function of these cell scores. For each gene, these models estimated the proportion of transcripts derived from cancer cells vs. immune and stromal cells in the average tumor (Supplementary Table 2)

Genes exhibited a wide range of cancer-derived expression (Figure 3A). Across all non-immune cancers, 5844 genes had less than 20% of transcripts attributed to cancer cells. Confirming the stability of this analysis, estimates of cancer-derived expression were largely consistent across TCGA datasets (Figure 3B). Confirming the specificity of this analysis, canonical marker genes were consistently estimated to have low percentages of transcripts from cancer cells. Gene lists used in many popular immune deconvolution algorithms^3,8,9,10,11,12^, most of which were designed for use in PBMCs and not in tumors, include substantial proportions of cancer-expressed genes (Figure 3C).

**Figure 3:**
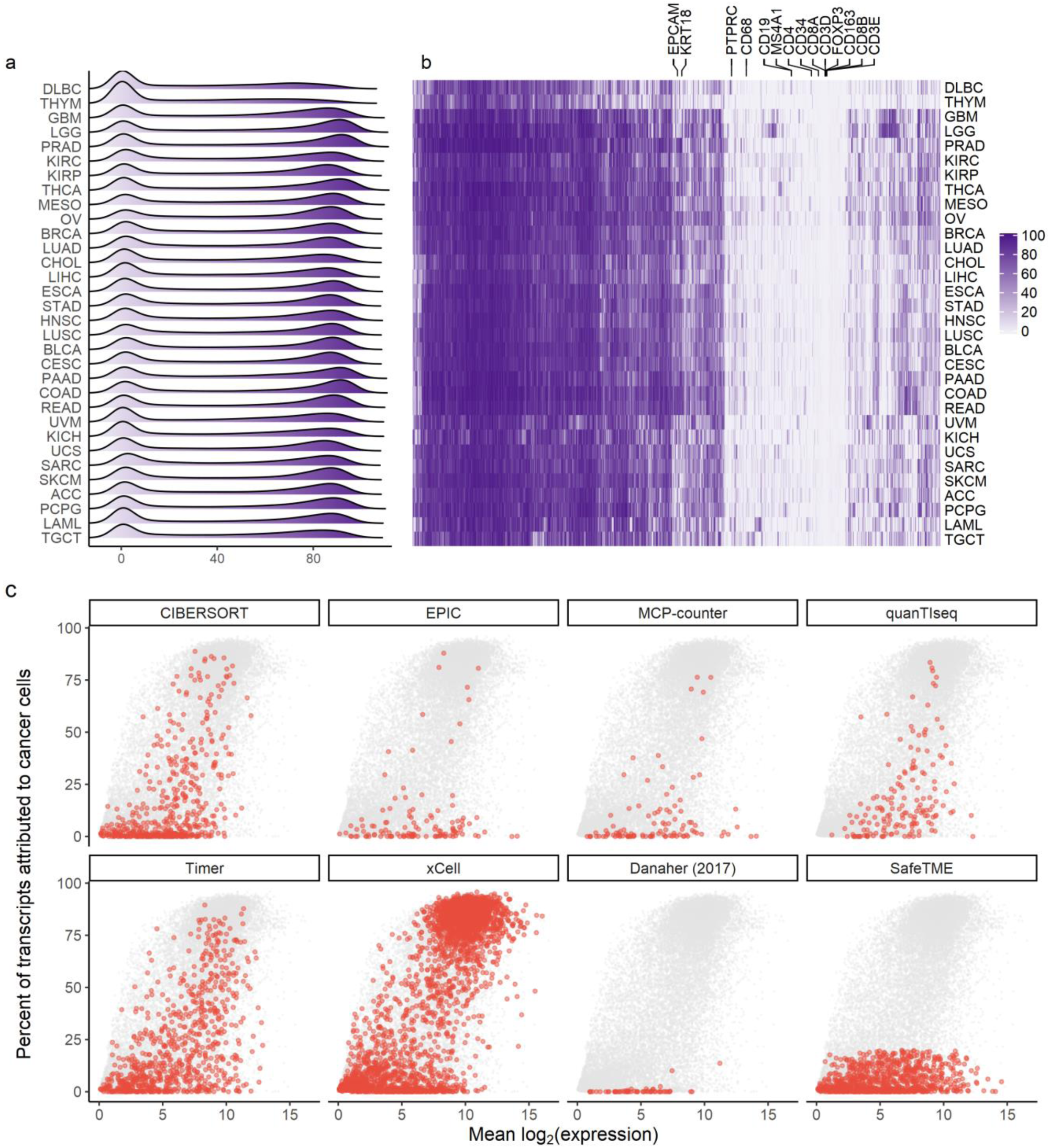
genes’ rates of cancer cell-derived vs. total expression in tumors. **a**. For each cancer type, density of genes’ percent of transcripts attributed to cancer cells. **b**. For all genes in all cancer types, estimated percent of transcripts attributed to cancer cells. **c**. Averaged across all non-immune tumors, genes’ mean expression vs. percent of transcripts attributed to cancer cells. Panels show gene lists from CIBERSORT, EPIC, MCP-counter, quanTIseq, Timer, xCELL, Danaher (2017), and SafeTME, the tumor-immune deconvolution cell profile matrix developed here.

### SafeTME: a cell profile matrix for deconvolution of the tumor microenvironment

To support deconvolution of the tumor microenvironment, we assembled the SafeTME matrix, a cell profile matrix for the immune and stromal cell types found in tumors. This matrix combines cell profiles derived from flow-sorted PBMCs^5^, scRNA-seq of tumors^13^ and RNA-seq of flow-sorted stromal cells^14^. It includes only genes estimated by the above pan-cancer analysis to have less than 20% of transcripts attributed to cancer cells.

### A library of scRNA-seq-derived cell profile matrices from diverse tissue types

To facilitate cell type deconvolution in diverse tissue types, we derived cell profile matrices from 27 publicly-available scRNA-seq datasets^15,16,17^ (Supplemental Table 3). In each dataset, we derived cell clusters, and we calculated the mean expression profile of each cluster. We named cell clusters using a combination of published marker genes^18^ and domain knowledge. Supplementary Table 4 details these cell profile matrices and the marker genes used to classify clusters.

### Harnessing the GeoMx platform to improve deconvolution’s capabilities

The GeoMx DSP platform extracts gene or protein expression readouts from precisely targeted regions of a tissue. First, the tissue is stained with up to four visualization markers, and a high-resolution image of the tissue is captured. Using this image, precisely-defined segments of the tissue can be selected for expression profiling; regions can be as small as a single cell or as large as a 700 µm ⨯ 800 µm region, and they can have arbitrarily complex boundaries. This flexibility in defining areas to be sampled is often used to split regions of a tissue into two segments, e.g. a PanCK^+^ cancer cell segment and a PanCK-microenvironment segment.

Two features of the GeoMx platform expand the abilities of mixed cell deconvolution. First, the platform counts the nuclei in every tissue segment it profiles. This nuclei count lets SpatialDecon estimate not just proportions but absolute counts of cell populations. The results of Figure 5 show cell population count estimates derived in this manner.

Second, the GeoMx platform can profile and model cell types that are absent in the pre-defined cell profile matrix. For example, when performing immune deconvolution in tumors, the expression profile of the cancer cells is often unknown. In such cases, the GeoMx platform can be used to select and profile regions of pure cancer cells, and this newly-derived cancer cell profile can be merged with the pre-defined cell profile matrix. This method is used to account for cancer cell expression in the deconvolution analyses of Figures 4 and 5.

**Figure 4:**
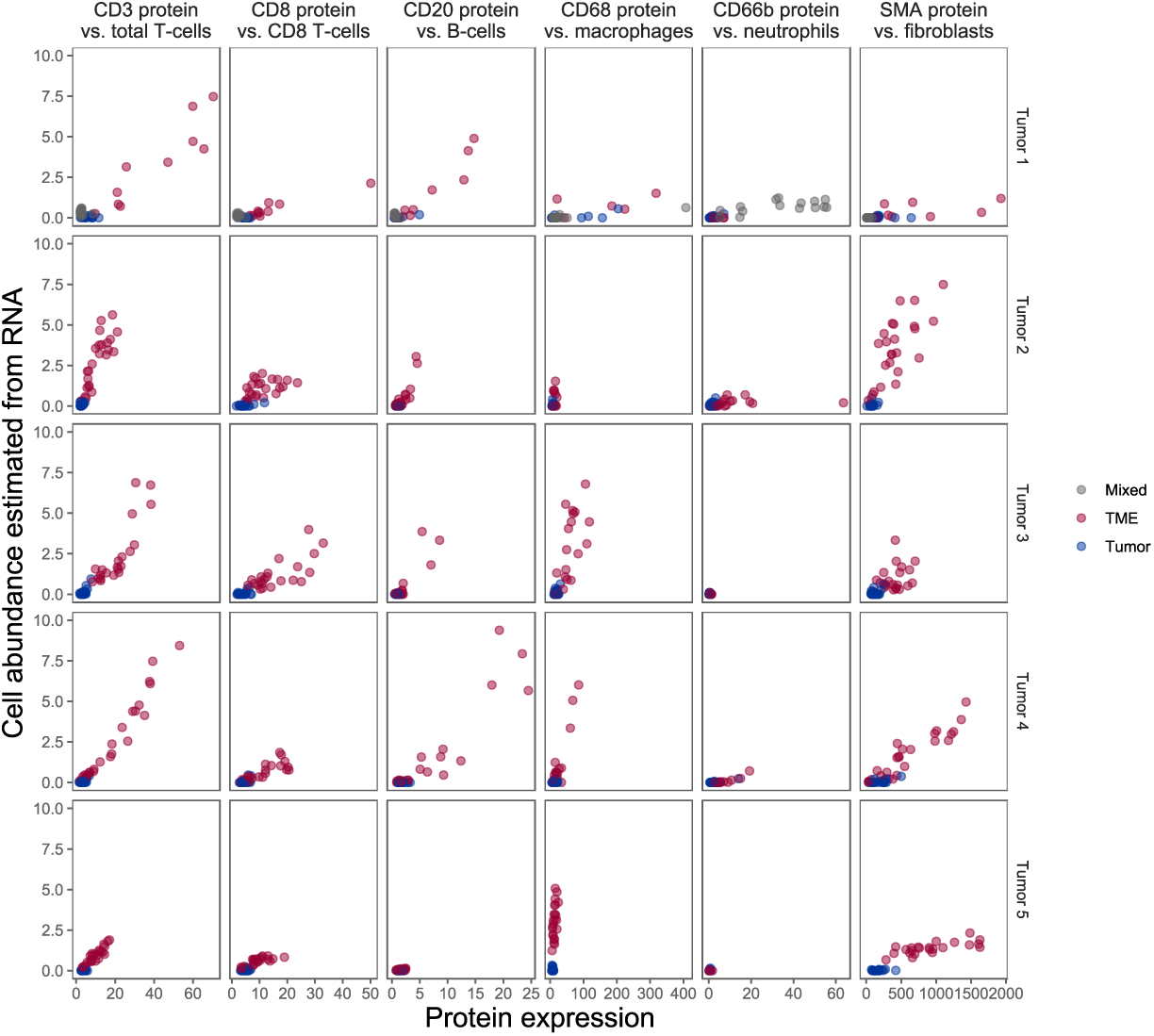
benchmarking of immune deconvolution vs. expression of canonical marker proteins. Each panel plots expression of a marker protein (horizontal axis) against a cell population’s abundance estimates from gene expression deconvolution (vertical axis). Tumor segments are in blue; microenvironment segments are in red. Each column of panels shows results from a single protein/cell pair; each row shows results from a different lung tumor.

**Figure 5:**
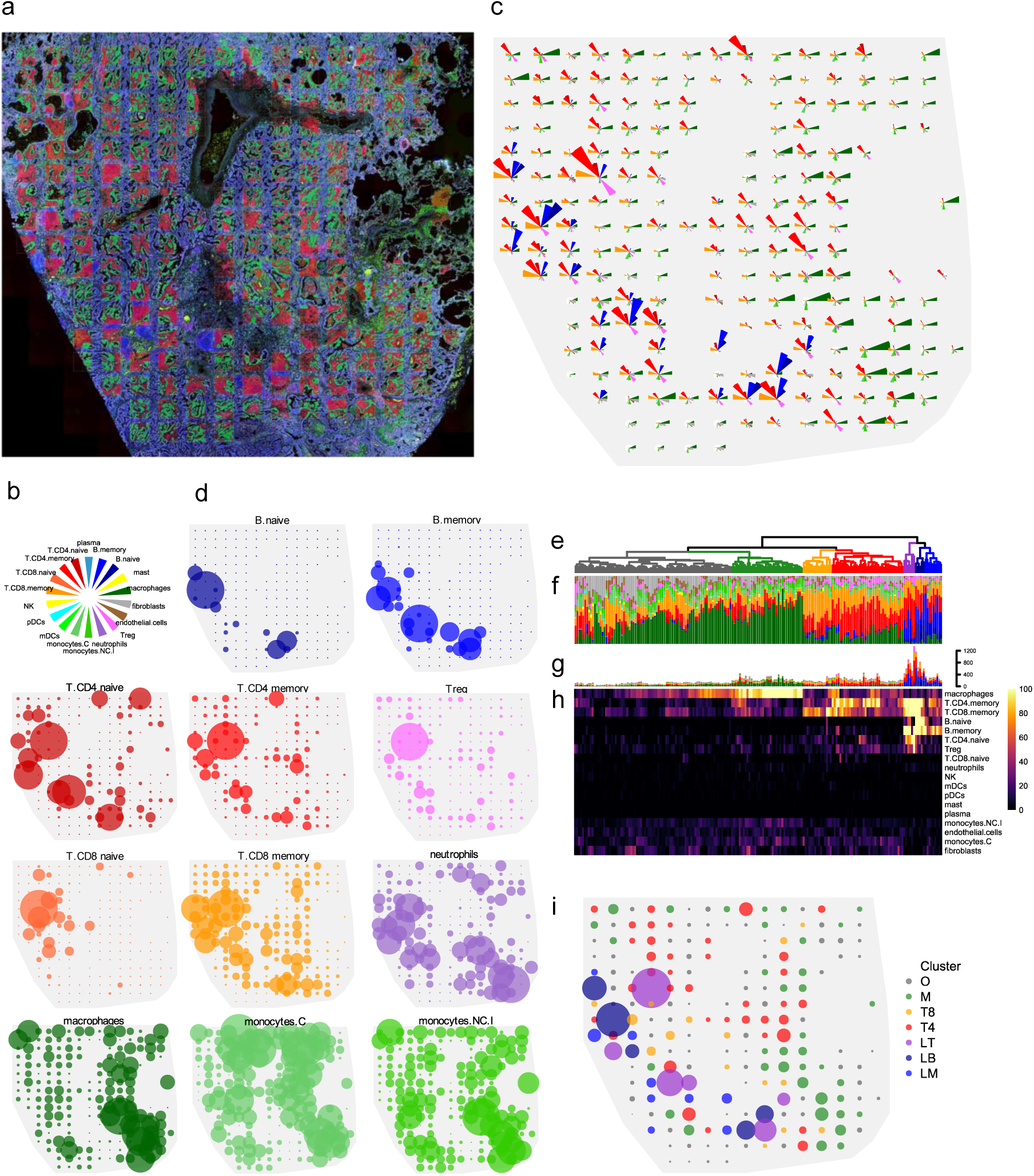
immune cell deconvolution in 191 microenvironment segments of a NSCLC tumor. **a**. Image of the tumor, with segments superimposed. Green = PanCK^+^ (tumor) segments; red = PanCK- (microenvironment) segments. **b**. Color key for panels c,d,f,g. **c**. Abundance estimates of 18 cell types in the microenvironment segments within 191 regions of the tumor. Wedge size is proportional to estimated cell counts. **d**. Abundance estimates of 12 cell populations in microenvironment segments. Point size is proportional to estimated cell counts within each panel; scale of point size is not consistent across panels. **e**. Dendrogram showing clustering of microenvironment segments’ abundance estimates. **f**. Proportions of cell populations in microenvironment segments. **g**,**h**. Estimated absolute numbers of cell populations in microenvironment segments. **i**. Spatial distribution of microenvironment segment clusters. Point color indicates cluster from (e); point size is proportional to total estimated immune and stromal cells in microenvironment segments.

### Paired spatially-resolved RNA and protein readouts provide a resource for benchmarking tumor-immune deconvolution methods in tissue

Due to practical limitations, most experiments benchmarking the performance of immune cell deconvolution methods rely on simulated data, generated either by in silico mixing of cell expression profiles^19^ or by in vitro mixing of purified cell populations^20^. However, simulations cannot faithfully represent performance in tumor samples: immune cell expression differs between blood and tumors^13^, and cancer cells can express putative immune genes. To benchmark deconvolution performance in real tumor samples, we used the GeoMx platform to collect paired measurements of gene expression and of canonical marker proteins.

From 5 FFPE lung tumors, we took two adjacent slides. We selected 48 700 µm regions from the first slide, and we identified their corresponding regions in the second slide. The selected regions in the first slide were profiled with the GeoMx protein assay, and the corresponding regions in the second slide were profiled with the GeoMx RNA assay, measuring 544 genes from the SafeTME matrix. Within each region, the GeoMx system’s flexible segmentation capabilities were used to collect separate profiles for tumor cells and for microenvironment cells.

To validate our algorithm, we compared deconvolution-derived cell estimates with the abundance of canonical marker proteins (Figure 4, Supplementary Table 5). In the average tissue, the correlation between protein expression and estimated cell abundance was 0.93 for CD3 protein vs. T-cells; 0.84 for CD8 protein vs. CD8 T-cells; 0.72 for CD68 protein vs. macrophages; 0.80 for CD20 protein vs. B-cells; and 0.80 for SMA protein vs. fibroblasts.

Neutrophils, whose low abundance in many tissues limited the range over which correlation could be observed, achieved an average correlation of just 0.43 with CD66b protein. However, in the two samples with the highest estimated neutrophils, this correlation rose to 0.86 (Tumor 1) and 0.84 (Tumor 4).

### Mapping the immune infiltrate in a NSCLC tumor

As a demonstration of spatially-resolved gene expression deconvolution, immune cell abundances were estimated across a grid of 191 regions of a NSCLC tumor. The GeoMx RNA assay was used to measure 1700 genes, including 544 genes from the SafeTME matrix. The tissue was stained with fluorescent markers for PanCK (tumor and epithelial cells), CD45 (immune cells), CD3e (T cells) and DNA. 191 300 ⨯ 300 µm regions of interest were arrayed in a grid across the 7.8 ⨯ 6.7 mm span of the tumor. Within each region of interest, the flexible illumination capability of the GeoMx platform was used to separately assay two segments: tumor segments, defined by PanCK^+^ stain, and microenvironment segments, defined as the tumor segments’ complement (Figure 5A).

The SpatialDecon algorithm was applied to all segments in the dataset using the SafeTME matrix along with tumor-specific profiles derived from the study’s PanCK^+^ segments. On a 1.9Ghz laptop, deconvoluting 376 segments took 29 seconds. Using just the microenvironment segments, we assem led a map of the tumor’s immune infiltrate (figure 5B). The most abundant cell types were CD8 T-cells (11,003 across all segments), CD4 T-cells (10,014), and macrophages (10,170). The algorithm estimated very low immune cell content in the tumor segments, with a mean of 7 immune and stromal cells per tumor segment, compared to a mean of 216 immune and stromal cells per microenvironment segment.

Cell populations had distinct spatial distributions. Naïve and Memory B-cells had the most concentrated spatial distributions (Gini coefficients = 0.84, 0.85), localizing primarily within a band of regions on the left side of the tumor. Naïve, memory and regulatory CD4 T-cell populations (Gini = 0.70, 0.64, and 0.57) had many dense foci near the B-cell-enriched regions and sporadic foci elsewhere in the tumor. Naïve CD8 T-cells (Gini = 0.52) concentrated in the top-right of the tumor, while memory CD8 cells were present throughout the tumor.

Macrophages (Gini = 0.48) and non-conventional/intermediate monocytes (Gini = 0.45, 0.39) were enriched in the lower-right of the tumor, away from the B-cells and T-cells, while conventional monocytes (Gini = 0.41) were enriched in the upper-right. Neutrophil-enriched segments (Gini = 0.41) appeared in both lymphoid-rich and myeloid-rich areas.

Hierarchical clustering on cell abundances identified 7 subtypes of tumor microenvironment regions (Fig 5e). The largest cluster, Subtype O, was defined by low total numbers of immune cells and consisted primarily of macrophages, memory CD8 T-cells, monocytes and fibroblasts. Subtype M was dominated by macrophages. Subtype T8 was dominated by memory CD8 T-cells, with less abundant memory CD4 T-cells. Subtype T4 was dominated by memory CD4 T-cells, with less abundant memory CD8 T-cells. Subtype LT consistent almost entirely of lymphoid cells, with majority T-cells but also abundant memory B-cells. Subtype LB also consistent almost entirely of lymphoid cells but had higher proportions of B-cells, both memory and naïve. Subtype LM was lymphoid-dominated but had as much as 15% macrophages. Each subtype was concentrated within, but not confined to, a distinct area of the tumor.

### Reverse deconvolution from cell abundances gives context to gene expression results

Variability in gene expression is driven both by changing abundance of cell populations and by differential regulation within cells. These two sources of variability can be decomposed via “reverse deconvolution”, in which each gene’s expression is predicted from cell abundance estimates. Outputs of this reverse deconvolution include genes’ fitted expression values based on cell abundances, and their residuals, calculated as the log2 ratio between observed and fitted expression (Fig 6a). These residuals measure genes’ up- or down-regulation within cells, independent of cell abundance.

**Figure 6:**
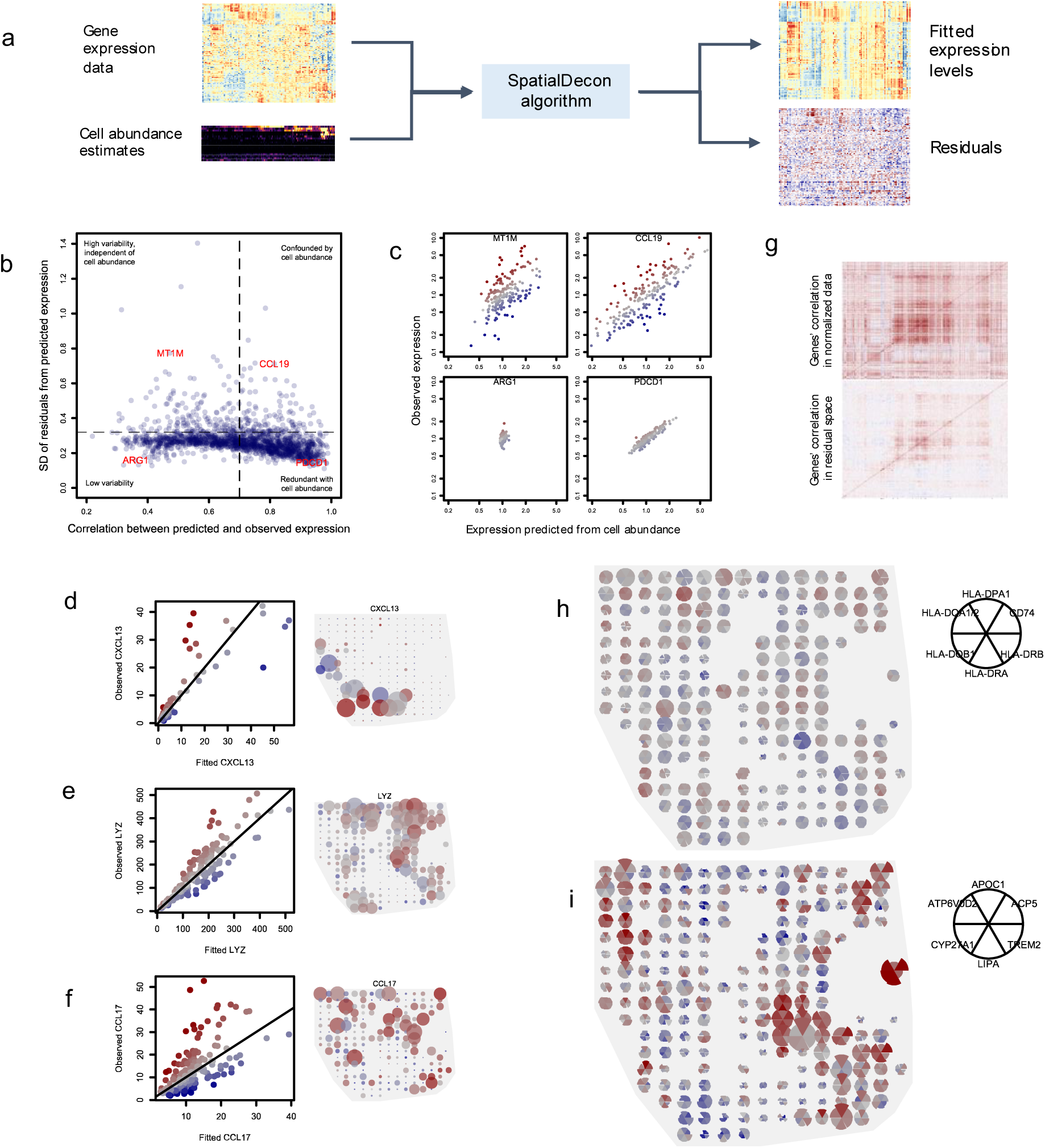
results of reverse deconvolution in a NSLCL tumor. **a**. Schematic of reverse deconvolution approach: gene expression is predicted from cell abundance estimates using the SpatialDecon algorithm, obtained fitted values and residuals. **b**. Genes’ dependence on cell mixing. Horizontal axis shows correlation between observed expression and fitted expression based on cell abundance. Vertical axis shows the standard deviation of the log2-scale residuals from the reverse deconvolution fit. **c**. Example genes from the extremes of the space of panel (b) are shown, with observed expression (vertical axis) plotted against fitted expression (horizontal axis). **d-f**. For CXCL13, LYZ and CCL17, observed expression is plotted against fitted expression (left), and observed expression is plotted in the space of the tissue (right). In all panels, point color indicates residuals. In panels on the right, point size is proportional to observed expression level. **g**. Correlation matrices of genes in log-scale normalized data (top) and in residual space (below). **h**,**i**. Spatial expression of gene clusters defined by high correlation in residuals of reverse deconvolution. Wedge color shows genes’ residual values; wedge size is proportional to genes’ expression levels.

To interpret our gene expression data in the face of highly variable cell mixing, we fit reverse deconvolution models over the microenvironment segments of the NSCLC tumor from Figure 5. Each gene’s dependency on cell mixing was measured with two metrics: the correlation between observed and fitted expression, and the standard deviation of the residuals. Based on these metrics, genes fell into 4 categories, each with a different implication for analysis and interpretation of genes’ data (Figure 6b). Genes with low correlations and high residual SDs, e.g. MT1M, are mostly independent of cell type mixing and can be understood without reference to cell abundances (Figure 6c). Genes with low correlations and low residual SDs, e.g. ARG1, have little variability to analyze. Genes with high correlations and low residual SDs, e.g. PDCD1, merely provide an obtuse readout of cell type abundance. Genes with high correlations and high residual SDs, e.g. CCL19, have substantial variability unexplained by cell mixing, but this variability is concealed by even greater variability driven by cell mixing. Analysis of these genes’ residuals reveals the full complexity of their behavior. For example, CXCL13 expression was over 2-fold higher or lower than expected in some regions (Figure 6d). LYZ expression, 84% of which was attributed to macrophages and monocytes, was highest in a corner of the tumor where those cell populations had relatively low abundance (Figure 6e). CCL17 was highly expressed in sporadic regions across the tumor, and in most of these regions the high expression was beyond what cell abundance alone could explain (Figure 6f).

### Correlation in residuals of reverse deconvolution reveals modules of co-regulated genes

Cell mixing induces correlation between genes that are expressed by the same cell type but that are not otherwise co-regulated. In the residuals of reverse deconvolution, this unwanted correlation abates, leaving only correlation induced by co-regulation (Fig 6g). For example, the correlation between CD8A and CD8B was 0.75 in the log2-scale data from microenvironment segments; in residual space, their correlation was -0.03. Correlation between MS4A1 and CD19 was 0.82 in the normalized data and 0.06 in residual space.

To identify candidate co-regulated genes, we identified gene clusters with high correlation in residual space. A cluster of CD74 and 5 HLA genes varied smoothly across the tissue, weakly correlated with macrophage abundance but also elevated in many macrophage-poor regions (Fig 6h). In the two regions with the most macrophages, these genes all had negative residuals, suggesting suppressed antigen presentation by macrophages in those regions. Another cluster consisted of lipid metabolism and small molecule transport genes (ACP5, APOC1, ATP6V0D2, CYP27A1, LIPA). Absolute expression of these genes was elevated in the tissue’s lower-right corner. Analysis of residuals reveals additional spatial expression dynamics, including a region of up-regulation in the upper-left side of the tissue and a region of down-regulation the lower-left (Fig 6i).

## Discussion

Cell deconvolution promises to be a linchpin of spatial gene expression analysis. Cell abundance estimates offer a functional significance and ease of interpretation unmatched by gene expression values. Cell abundance also gives context to gene expression results, disam biguating whether a gene’s expression pattern results from differential cell type abundance or differential expression within cell types.

The methods described here enable spatial studies as a natural follow-on to scRNA-seq: given cell populations defined by scRNA-seq, deconvolution in spatial gene expression data reveals how those cells are arranged within tissues, obtaining a region-by-region accounting of their abundance. This allows new questions to be asked: How are cell types arranged and mixed with each other? Which cell types repel or attract each other? Which cell types explain the expression pattern of a gene of interest? How does a cell population’s behavior change when it is co-localized with another cell population?

The methods and data resources described here promise to improve deconvolution not just in spatial expression data but also in bulk gene expression. Log-normal regression has the same theoretical benefits in bulk expression deconvolution. Our library of cell profile matrices for diverse tissues directly supports deconvolution in bulk gene expression experiments. And future attempts to deconvolve immune cells in bulk tumor expression data should confine analysis to our list of genes not expressed by cancer cells.

Based on cell abundances, we identified 7 microenvironment subtypes within one NSCLC tumor. This heterogeneity raises the prospect that tumors could be classified not just by their overall cell abundance, but by the localized microenvironment subtypes they contain.

SpatialDecon, an R library implementing these methods, is available at https://github.com/Nanostring-Biostats/SpatialDecon. The library of cell profile matrices is available at https://github.com/Nanostring-Biostats/CellProfileLibrary. The in-situ benchmarking dataset is available at https://github.com/Nanostring-Biostats/ImmuneDeconBenchmark.

## Methods

### The SpatialDecon algorithm

Notation

Let X_p.K_ be the cell profile matrix giving the linear-scale expression of p genes over K cell types.

Let Y_p.n_ be the observed expression matrix of p genes over n observations.

Let β_K.n_ be the unobserved matrix of cell type abundances of K cell types over n observations.

Let B_p.n_ be the matrix of expsubject to the constraintected background counts corresponding to each data point in Y.

Let ||x|| denote the L2 norm operator of x such that ||x|| = mean(x^2^).

The core log-normal deconvolution algorithm proceeds as follows:

#### Algorithm 1

1. To avoid negative-infinity values when log-transforming Y, define ε equal to the minimum non-ero value in Y, and threshold Y elow so that its smallest value is ε.
2. Take 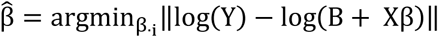, subject to the constraint that β._*i*_≥ 0. This constrained optimization is performed separately for each column of Y using the R package logNormReg^6^.
3. For i in {1,…, n}, calculate the covariance matrix of by 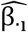 inverting the hessian matrix returned by logNormReg. Call this covariance matrix 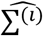. Then the standard error for 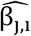 is 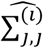.
4. Calculate the p-value for each β_i,j_ with 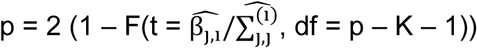, where F is the cumulative distribution function of the t distribution.

The SpatialDecon algorithm, which incorporates outlier removal into algorithm 1, proceeds as follows:

#### Algorithm 2 (SpatialDecon)

1. Run Algorithm 1.
2. Choose δ as the expression level elow which technical noise predominates. For GeoMx data normalized to have expected background = 1, we use δ = 0.5.
3. Define the residuals of the algorithm fit as 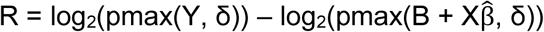, where pmax(x, δ) is the function replacing all elements of x elow δ with δ.
4. For all {i,j} with |R_i,j_| > 3, set Y_i,j_ to NA.
5. Re-run Algorithm 1 using the updated Y matrix.

### Using the GeoMx platform to derive cell profiles for new cell types

The procedure for using GeoMx to derive the profiles of unmodelled cells and merge them into the cell profile matrix X proceeds as follows:

#### Algorithm 3

1. Specify columns of Y corresponding to segments selected to contain a pure cell type that is missing from X. For example, for immune deconvolution in tumors, select segments targeting purely PanCK^+^ cells to derive a cancer cell profile.
2. Collapse the segments into 10 clusters by applying the R functions hclust and cutree to their log-transformed expression profiles.
3. Define each cluster’s expression profile by taking each gene’s geometric mean across the observations in the cluster. Scale this profile so its 90^th^ percentile value is equal to the average 90^th^ percentile value of the columns of the cell profile matrix.
4. Append the cluster expression profiles to the cell profile matrix.

The NanoString GeoMx® Digital Spatial Profiler and GeoMx assays are for research use only and not for use in diagnostic procedures.

### Using the GeoMx platform to convert cell abundance scores to cell counts

When the GeoMx system’s per-region nuclei counts are available, the below procedures are used to convert cell abundance scores to estimates of absolute cell counts.

Case 1: all cell types in the tissue are modelled in the cell profile matrix: Here we estimate the number of each cell type in a region by the product of the nucleus count in the region and the cell type’s estimated proportion in the region: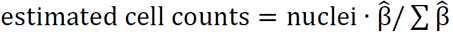.

Case 2: the tissue contains cell types that are not modelled by the cell profile matrix. The motivating case here is immune cell deconvolution in tumors, where cancer cell profiles are often omitted from the model. If it is reasonable to assume that at least one profiled region consists of entirely cells modelled by the cell profile matrix, then call the sum of its cell a undance scores β_max_. Then for all regions, take estimated cell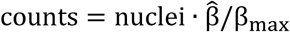.

### Analysis of cell pellet array study

Reference genes for normalization were selected by applying the geNorm algorithm^21^ to the 50 highest-expressing genes. Each segment’s expression profile was normalized using the geometric mean of the resulting list of 27 reference genes. The expression profiles of the pure cell lines were estimated using the median expression profile of the 4 unmixed replicates from each cell line. These two profiles were then scaled to have the same median expression level.

Gene sets were defined as follows. The “most informative” gene set had a log2 fold-change between the cell lines of > 1.5, for a total of 1 3 genes. The “least informative” gene set had a log2 fold-change < 1.5, for 1231 genes. The “complete” gene set was 1414 genes, and the “low-to-moderate expression” genes had <1000 counts in each cell line, for 1362 genes.

Log-normal deconvolution was run using Algorithm 1. Non-negative least squared deconvolution was run by taking 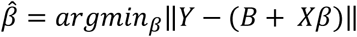, subject to the constraint that β ≥ 0. Optimization was performed using the R function optim. The background term B was included because ignoring background would disadvantage NNLS in the comparison. Nu support vector regression was run using svm function from package e1071, with v set to 0.75, a linear kernel, and without scaling. v-SVR does not allow for explicit modelling of background signal, so normalized expression data was background-subtracted before entry into v-SVR.

To compute each gene’s influence on a deconvolution result, deconvolution was run once with the complete gene set and once with each gene omitted. Each gene’s influence was reported as the absolute value difference in estimated HEK293T proportion between deconvolution with the complete gene set and deconvolution with the leave-one-out gene set.

### Analysis of TCGA for identifying genes suitable for use in immune deconvolution in tumor samples

Each TCGA sample was scored for abundance of diverse immune and stromal cells using the geometric mean of previously reported marker genes^7,8^. Then, in each cancer type, we used lognormal regression to model each gene as follows:

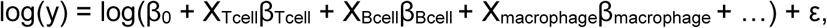

where y is the vector of the gene’s (linear-scale) expression across all samples in a cancer type, X_cell_ is the vector of a cell type’s estimated abundance across all samples, and ε is a vector of normally-distributed noise. We additionally apply the constrains that all β terms are ≥ 0. The use of lognormal regression is motivated by the same considerations used in our deconvolution method: expression from mixed cell types compounds additively, but noise in gene expression is a log-scale phenomenon.

In this model, the β_0_ term represents the gene’s average expression in a tumor when no immune cells are present. Then we can measure a gene’s proportion of tumor-intrinsic expression with β_0_ / mean(y). Ideal genes for deconvolution will have β_0_ / mean(y) very close to 0; genes with substantial contamination from cancer cells will have β_0_ / mean(y) near 1.

### Derivation of the SafeTME cell profile matrix for deconvolution in the tumor microenvironment

Three datasets were used to define the SafeTME cell profile matrix for deconvolution of the tumor microenvironment: expression profiles of flow-sorted PBMCs for use in deconvolution of blood samples^5^, scRNA-seq of finely clustered immune cell types^13^, and RNAseq profiles of 6 cell populations flow-sorted from lung tumors^14^.

Cell type profiles from PBMCs were used whenever possible, since flow-sorting on surface markers is the gold standard for classifying immune cells. Specifically, profiles were taken for naïve B cells, memory B cells, plasmablasts, naïve CD4 T-cells, memory CD4 T-cells, naïve CD8 T-cells, memory CD8 T-cells, T-regulatory cells, NK cells, plasmacytoid DCs, myeloid DCs, conventional monocytes, non-conventional/intermediate monocytes, and neutrophils. We omitted profiles of PMBC cell populations expected to be vanishingly infrequent in tumors: basophils, MAIT cells, and T gamma delta cells.

From the tumor scRNA-seq dataset, we took the profiles for macrophages and mast cells, which are not present in PBMCs. We defined a mast cell profile as the average of the 2 reported mast cell clusters’ profiles, and the macrophage profile as the average of the 9 reported macrophage cluster profiles^13^. The mast cell profile was scaled to have the same 80^th^ percentile and the average 80^th^ percentile of the PMBC cell profiles; the macrophage profile was scaled to have the same 80^th^ percentile as the PBMC conventional monocytes profile.

From the flow-sorted lung tumor dataset^14^, we derived profiles for endothelial cells and fibroblasts. Four endothelial cell samples with low signal were removed, as were 8 fibroblast samples with low signal. The remaining replicate samples were normalized using their 90^th^ percentiles, and endothelial cell and fibroblast profiles were defined by the median expression profiles of the corresponding samples. The cell profiles extracted from the 3 datasets then were combined into a single matrix, which was reduced to a subset of 1180 highly-informative genes.

It is inevitable that the combined matrix contains numerous systematic biases, such as platform effects, noise in the experimental results of the original cell profile matrices, and gene expression differences in blood vs. tumor. To reduce these effects, we employed the following procedure. First, we performed deconvolution on 3 TCGA datasets: colon adenocarcinoma (COAD), lung adenocarcinoma (LUAD), and melanoma (SKCM). Most genes were consistently over- or under-estimated by the deconvolution fits, and these biases were consistent across datasets. Each gene’s bias was estimated with the geometric mean of the ratios between its observed expression values and its predicted expression values from the deconvolution fits.

Finally, each gene’s row in the cell profile matrix was then rescaled by its expected bias.

We removed genes estimated by our TCGA analysis to have more than 20% of transcripts derived from tumor cells. The final SafeTME cell profile matrix, from 18 cell types and 906 genes, is reported in the Supplementary data.

### Derivation of cell profile matrices from public scRNA-seq datasets

27 single cell RNA-seq studies were downloaded from The Broad Institute Single Cell Portal. For raw gene expression (GE) matrices, cells were removed if they fell below the inflection point, were considered empty by emptyDrops^22^, had a gene count above 2.5x average gene count, or had a percentage of mitochondrial genes > 0.05. Genes were removed if they appeared in less than 2 cells or had low biological significance as measured by scran^23^. Cells were clustered and marker genes identified using Seurat^24^. Clustered marker genes were compared to PanglaoDB^18^ marker genes (ubiquitousness index < 0.1, sensitivity > 0.6, specificity < 0.4, canonical marker). Cell clusters were named according to the PanglaoDB cell type with the most overlapping marker genes. All cell cluster names were manually reviewed for correctness. When data sets had already been annotated with cell type calls, the existing cell type calls were retained. Only cell clusters with more than 10 cells were reported. Each cell cluster’s profile was reported as the arithmetic mean of its cells’ expression profiles.

### Protein Slide Preparation

For GeoMx DSP slide preparation, we followed GeoMx DSP slide prep user manual (MAN-10087-04). 5 µm FFPE microtome sections of non-small-cell-lung cancers (NSCLC) (ProteoGenex) or cell pellet arrays (Acepix Biosciences, Inc.) were mounted onto SuperFrost Plus slides (Fisher Scientific, 12-550-15) and air dried overnight. Slides were prepared by baking in a drying oven at 60°C for 1 hour; then the paraffin was removed with CitriSolv (Fisher Scientific, 04-355-121). The samples were rehydrated in an ethanol gradient and final wash in DEPC-treated water (ThermoFisher, AM992). Target retrieval was performed by placing slides in staining jars containing 1x citrate buffer pH6 (Sigma Aldrich SKU C9999-1000ML) and heated in a pressure cooker on high temperature setting for 15 minutes. Slides were allowed to cool to room temperature and blocked at room temperature for one hour with Buffer W (NanoString Technologies). The primary antibody mix was made by combining the detection antibody modules (NanoString Technologies) at 1:25 and the visualization markers in Buffer W. The NSCLC were visualized with CD3-647 at 1:400 (Abcam, ab196147), CD45-594 at 1:40 (NanoString Technologies) and PanCK-532 at 1:40 (NanoString Technologies. Slides were incubated overnight at 4°C. Slides were fixed with 4% paraformaldehyde (Thermo Scientific 28908) and the nuclei were stained with SYTO 13 (Thermo Scientific S7575) at 1:10 for 15 minutes.

### RNA/NGS Slide Preparation

For GeoMx DSP slide preparation, we followed GeoMx DSP slide prep user manual (MAN-10087-04). 5 µm FFPE microtome sections of both non-small-cell-lung cancers (NSCLC) (ProteoGenex) were mounted onto SuperFrost Plus slides (Fisher Scientific, 12-550-15) and air dried overnight. Slides were prepared by baking in a drying oven at 60°C for 1 hour. Slides were then processed with a Leica Biosystems BOND RXm (Leica Biosystems) as specified by the NanoString GeoMx DSP Slide Preparation User Manual (NanoString Technologies, MAN-100 7). Briefly, slides were processed with the taining protocol “*GeoMx RNA DSP slide prep”, the Preparation protocol “* Bake and Dewax”, HIER protocol “*HIER 20 min with ER2 @ 100°C, and Enzyme protocol “*Enzyme 1 for 15 minutes”. For Enzyme 1 a 1 ug/mL concentration of Proteinase K (Ambion, 2546) was used. This program included target retrieval, Proteinase K digestion, and post fixation. Once the Leica run had finished slides were immediate removed and placed in 1x PBS. One at a time, slides were placed in a prepared HybEZ Slide Rack in a HybEZ Humidity Control Tray (ADC Bio, 310012) with Kimwipes damped with 2xSSC lining the bottom. 200uL of a custom RNA probe Mix at a concentration of 4nM per probe in 1x Buffer R (NanoString Technologies), was applied to each slide. A Hybridslip (Grace Biolabs, 714022) was immediate applied over each sample. Slides were incubated in a HybEZ over (ACDBio 321720) at 37°C for 16-24 hours. After hybridization slides were briefly dipped into a 2x SSC + 0.1% Tween-20 (Teknova, T0710) to allow the coverslips to slide off then washed twice into a 2x SSC/50% formamide (ThermoFisher AM9342) solution at 37°C for 25 minutes each, followed by two washes in 2x SSC for 5 minutes each at room temperature. Slides were then blocked in Buffer W (NanoString Technologies) at room temperature for 30 minutes. 200uL of a morphology marker mix was them applied to each sample for 1 hour. The tumors were visualized with CD3-647 at 1:400 (Abcam, ab196147), CD45-594 at 1:10 (NanoString Technologies), PanCK-532 at 1:20 (NanoString Technologies) and SYTO 13 at 1:10 (Thermo Scientific S7575).

### GeoMx DSP sample collection

For GeoMx DSP sample collection, we followed GeoMx DSP instrument user manual (MAN-10088-03). Briefly, tissue slides were loaded to GeoMx DSP instrument and then scanned to visualize whole tissue images. For cell pellet array samples, 300um ROIs in diameter were placed. For each tissue sample, we placed ROIs and segmented into two regions: PanCK-high tumor region and PanCK-low TME regions.

### GeoMx DSP NGS Library Preparation and Sequencing

Each GeoMx P sample was uniquely indexed using Illumina’s i5 ⨯ i7 dual-indexing system. 4 uL of a GeoMx DSP sample was used in a PCR reaction with 1 uM of i5 primer, 1 uM i7 primer, and 1X NSTG PCR Master Mix. Thermocycler conditions were 37°C for 30 min, 50°C for 10 min, 95°C for 3 min, 18 cycles of 95°C for 15 sec, 65°C for 60 sec, 68°C for 30 sec, and final extension of 68°C for 5 min. PCR reactions were purified with two rounds of AMPure XP beads (Beckman Coulter) at 1.2x bead-to-sample ratio. Libraries were paired-end sequenced (2×75) on a NextSeq550 up to 400M total aligned reads.

### Analysis of GeoMx protein and RNA benchmarking data

Twelve segments with very low signal in either the protein or RNA results were excluded. The protein assay data was normalized with the geometric mean of the negative control antibodies, and the RNA data was normalized with the geometric mean of the negative control probes. Prior to deconvolution, Algorithm 3 was used to append tumor-specific profiles to the SafeTME matrix. Deconvolution was run using the resulting profile matrix and the SpatialDecon algorithm.

### Analysis of GeoMx RNA data from a grid over a NSCLC tumor

Raw counts from each gene in each tissue region were extracted from the NanoString GeoMx NGS processing pipeline. For each region, the expected background for each gene was estimated with the mean of the panel’s 100 negative control probes. Each region’s signal strength was measured with the 85^th^ percentile of its expression vector. Three PanCK^+^ regions with outlier low signal strength were removed from the analysis. Each region’s data was normalized with the “signal-to-background” method, scaling each region such that its negative control probe mean was 1. (This method is one of the manufacturer’s recommended approaches for normalizing GeoMx data; it is successful because the negative control probes respond to technical factors like region size, region-specific RNA binding efficiency, and region-specific density of material to which oligos might bind.)

Prior to deconvolution, the study’s pure PanCK^+^/tumor segments were input into Algorithm 3, resulting in 10 tumor-specific expression profiles. These profiles were then appended to the SafeTME profile matrix. Deconvolution was then run using the SpatialDecon algorithm.

To derive microenvironment subtypes, clustering was performed on the matrix of estimated cell counts using the R library pheatmap.

### Methods: calculation of residuals from cell scores in NSCLC study

Reverse deconvolution was run as follows. Cell abundance estimates were taken from the SpatialDecon run described above. In only the stroma segments, each gene’s linear-scale expression was predicted from the cell abundance estimates, including an intercept term. All estimates were constrained to be non-negative.

Reverse deconvolution residuals were calculated for each gene as the log2 fold-change between observed expression and fitted expression, with both terms thresholded below at 1, the expected background in the normalized data. That is, if y.observed is a gene’s normalized expression, and y.fitted is its predicted expression based on the reverse deconvolution fit, then residuals = log2(max(y.observed, 1)) – log2(max(y.fitted, 1)).

The following metrics were used to measure genes’ dependency on cell mixing. “Correlation” was calculated as cor(y.observed, y.fitted). “Residual “was calculated as sd(residuals).

To identify clusters of co-regulated genes, the correlation matrix of all genes’ reverse deconvolution residuals was clustered using the R function hclust. Gene modules were identified by applying the R function cutree to the resulting hierarchical clustering results.

## List of supplementary files

Supplemental Table 1: Marker genes used in pan-cancer screen for genes with minimal cancer-intrinsic expression.

Supplemental Table 2: Percent of transcripts attributed to cancer cells for each gene in TCGA.

Supplemental Table 3: Details of the library of cell profile matrices for deconvolution of diverse tissue types.

Supplemental Table 4: Marker genes for the cell types in the library of cell profile matrices.

Supplementary Table 5: Correlations between canonical marker proteins and deconvolution results, for each cell type and tissue in Figure 4.

## Supplementary material

**Supplementary results: lognormal regression is a more appropriate model for mixed cell expression data**

Least-squares-based deconvolution algorithms solve extensions of the optimization problem ||y – Xβ||, where y_p.1_ is a sample’s (linear-scale) expression levels of p genes, X_p.K_ is the cell profile matrix containing the expected expression of p genes in K cell types, and β_K.1_ is the vector of the K cell types’ abundances, and ||*|| is the L2 norm operator.

Implicit in any deconvolution technique is a mean model, specifying how cell types’ expression profiles add up to create a mixed profile; and a variance model, describing the noise between expected and observed expression. Least squares regression’s mean model assumes E(y) = Xβ. This mean model is appropriate, as the gene expression profiles of different cell types should add atop each other in a mixed sample. But least squares regression’s variance model is inconsistent with the gene expression data.

Least squares regression’s variance model assumes that every gene’s residuals (y – Xβ) are normally distributed with equal variance. This model is inappropriate for gene expression data for two reasons. First, gene expression data is right-skewed, strongly violating the assumption of normality (Supp. Fig. 1 a,b). Second, gene expression data is heteroscedastic, with high-expressing genes having standard deviations thousands of folds above low expressing genes (Supp. Fig. 1 c). Log-transforming gene expression data largely corrects both its skew and its heteroscedasticity.

Supplementary Figure 1 uses data from the TCGA LUAD (lung adenocarcinoma) RNAseq dataset to demonstrate gene expression’s skewness and heteroscedasticity on the linear-scale and its relative normality and homoscedasticity on the log-scale. Panel (a) shows the distribution of CD274 (PD-L1), a gene with typical skewness. On the linear scale its skewness is 2.8; after log-transformation its skewness is 0.1 (the normal distribution has skewness of 0). Panel (b) shows the distribution of the skewness statistics from all genes in the transcriptome in TCGA LUAD. On the linear-scale, all but one of the 20243 genes measured was right-skewed, and 68% have extreme skewness > 2. On the log-scale, the average gene has skewness close to 0, and only 0.4% of genes have skewness outside of (−2, 2). Panel (c) demonstrates the heteroscedasticity of gene expression, plotting each gene’s SD against its mean. In linear-scale data, SD increases proportional to a gene’s mean expression level, and the range of SDs spans 9.6.10^−3^ to 1.9.10^5^ (20004870-fold). In log-scale data, low-expression genes are only slightly more variable than high-expressors, and the range of SDs is only 16.5-fold.

**Supplementary Figure 1:**
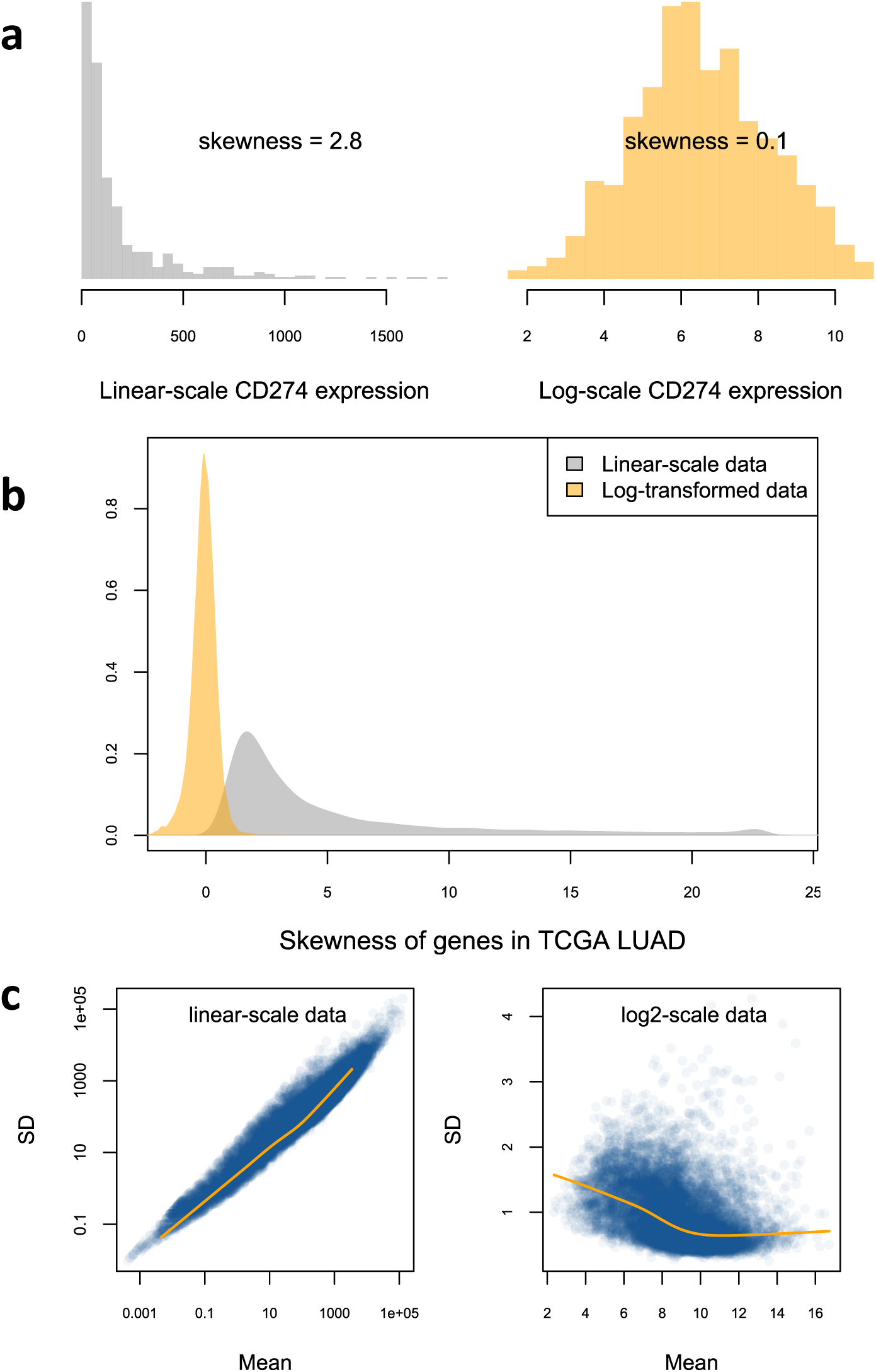
skewness and heteroscedasticity of gene expression data. All figures are generated from the TCGA LUAD dataset. **a**. Histograms of CD274 (PD-L1) expression on the log and linear scale. **b**. Distribution of skewness statistics calculated for each of 20531 genes across the TCGA LUAD samples. Grey: skewness of linear-scale genes. Orange: skewness of log-scale genes. **c**. Genes’ mean vs. standard deviation, calculated from linear-scale and from log-scale data.

**Supplementary Figure 2:**
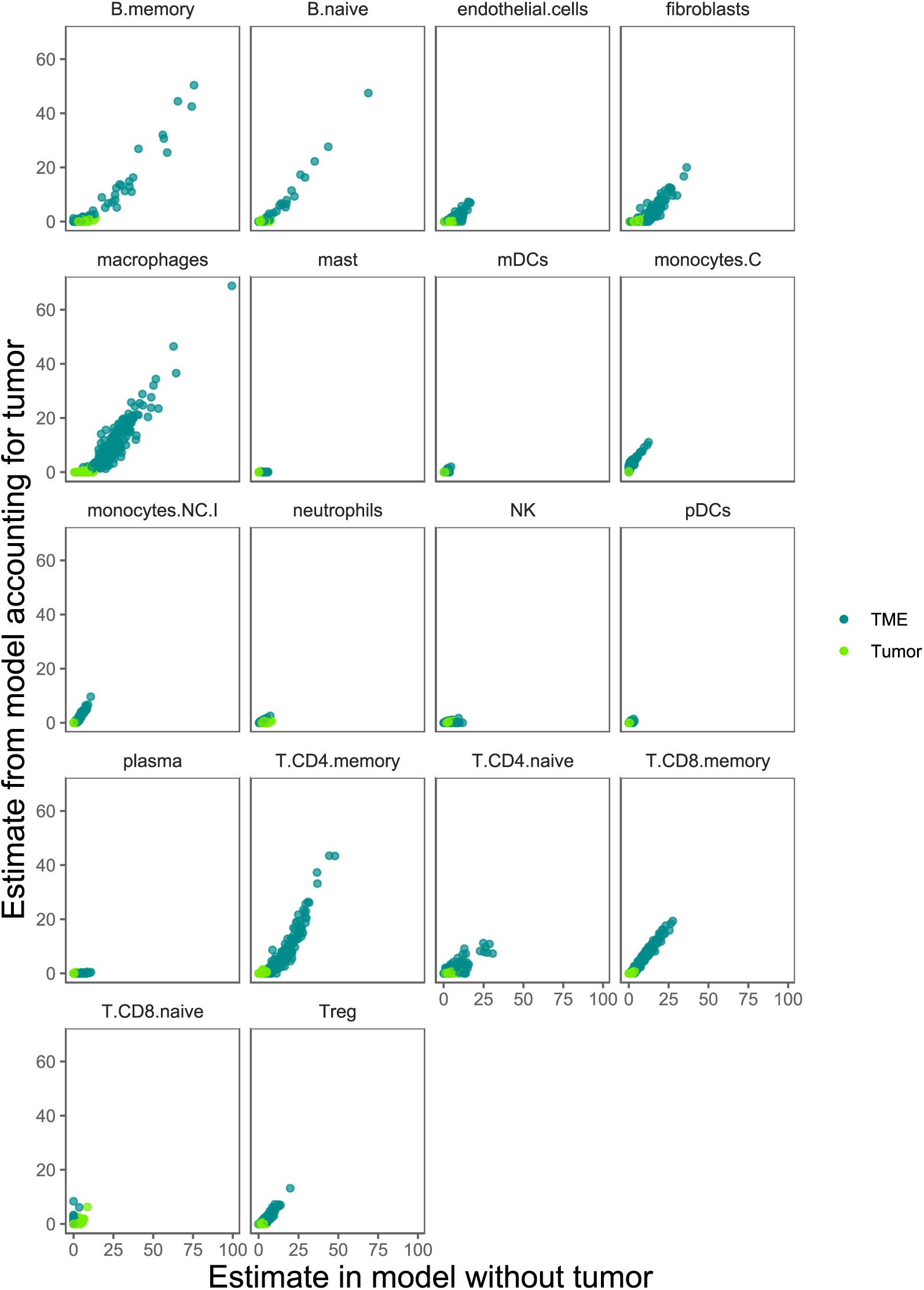
Stroma cell abundance estimates from segments of a NSCLC tumor, with and without modelling tumor-specific expression. **Horizontal axis:** Estimates from deconvolution using only stroma cell profiles. **Vertical axis:** Estimated from deconvolution using both stroma cell profiles and 10 tumor cell profiles derived from pure tumor segments.

